# The flow of sickle blood through glass capillaries: a point of care application

**DOI:** 10.1101/837369

**Authors:** Christopher D. Brown, Alexey M. Aprelev, Frank A. Ferrone

## Abstract

We show that the characteristic time for sickled blood to traverse 100 µm-diameter glass capillaries can be used as the foundation for a useful point-of-care diagnosis for sickle cell disease. 3.2 cm long capillaries were inserted into a 5 µL drop of blood, and 1/4 µL was spontaneously drawn into the tube. The rate at which the blood traversed the capillary was significantly changed by deoxygenation only for sickle genotypes, a consequence of the increased viscosity of cells containing sickle fibers. Under these conditions, we find deoxygenation causes the time for sickle blood to traverse the capillary to increase to about 3 times its original value, in contrast to normal blood, where the times are equivalent regardless of state of oxygenation. This corresponds to readily-observed time differences of around 10 s for capillary traversal. Such changes are present for blood from homozygous sickle cell patients, as well as heterozygous patients with hemoglobin AS, SC, and Sβ^+^ thalassemia. Moreover, we demonstrate that this test is sensitive to sickling therapies that increase the production of fetal Hb. This viscosity-based measurement thus offers the prospect of an assay that is 10 times faster than any current test of which we are aware, at costs that rival the least expensive of any current assays.

## Introduction

The life expectancy of a child born with Sickle Cell Anemia is drastically dependent on where that child is born. Were the child born in the US, he or she can expect a life-span of around 40 years, on average, [1] but for the same child born in Africa, it is under 5 years. [2] [3] Alas, 79% of sickle cell newborns are found in sub-Saharan Africa, and by 2050 this is expected to rise to a phenomenal 88%.[4] The overall standard of medical care will undoubtedly play a role in any lifespan, to be sure, but administration of prophylactic penicillin [5] and pneumococcal vaccination [6] are known to have a profound effect on the life expectancy of sickle cell patients in developing nations, once the children at risk are identified [3]. The tests typically used in the developed world to make this determination, such as isoelectric focusing, or HPLC, not only are prohibitively expensive, but also tend to require highly trained personnel. This makes a simple and rapid test an important goal for public health, especially in regions of high incidence. Indeed, the recognition of the value of a rapid diagnostic test was first realized by no less than Linus Pauling some 70 years ago.[7]

Sickle Cell Anemia is the most severe of the sickle cell genotypes, since the patient is homozygous for HbS. A variety of heterozygous conditions exist. The most common is to mix one gene for S with one for normal HbA. The AS combination is known as sickle trait, and is largely benign. In Sβ^+^ thalassemia, one again has a combination of A and S, but with far less A, and patients with Sβ^+^ thalassemia suffer from many of the same issues as SS patients [8, 9]. Finally, combining an S gene with one for HbC produces a phenotype, SC, with severity comparable to that of Sβ^+^ thalassemia [8, 9].

The damage inflicted by sickle cells, of any composition, begins when deoxygenation rigidifies the mass of hemoglobin they contain. [10, 11] By obstructing the microcirculation, these cells deprive downstream tissues of oxygen, producing organ damage, and painful episodes. Chien et al [13] demonstrated that chemically hardening of the red-cell membrane of normal blood cells by acetaldehyde to make erythrocytes undeformable elevated the viscosity several times, and thus it is unsurprising that cells that cannot deform because of internal rigidity likewise display elevated viscosity, as has been observed for some time.[12] We hypothesized that this induced stiffness could be used as a simple and rapid diagnostic for the presence of the sickle gene. The needs of economy and portability rule out conventional viscometers, but the flow of blood into a small glass capillary might have the requisite properties. We thus were led to explore the flow of whole blood drawn into a 100 µm-diameter capillary, comparing deoxy blood to its oxygenated counterpart.

We find that this simple measurement has great potential, and rests on a solid foundation as shown elsewhere. Sickled (i.e. rigidified) cells flowed through these capillaries significantly more slowly than their flexible counterparts. While this flow can be described as the behavior of a simple fluid with elevated viscosity for much of the flow, the rigid cells were also found to accumulate into a dense zone or cap behind the advancing meniscus, retarding the velocity even further. Although these features appeared in all instances of sickle cell disease, we also observed systematic differences between the different genotypes. In this paper we explore the common features as well as the systematic differences, and show to what extent clinically useful discrimination is possible. The data collected here used cameras and positioning equipment for detailed characterization, but, as we show, visual observation and a simple stopwatch are all that is required for an efficient and inexpensive assay.

## Materials and Methods

Blood was collected from 23 patients, primarily from St. Christopher’s Hospital in Philadelphia. In addition, AA blood was contributed by the investigators, while AS blood was contributed from the laboratory of Dr. W. A. Eaton of the NIH as a discard product from a study underway there. Blood was collected into EDTA anticoagulant in all cases. A 5 µL drop was used for the measurements, and in the case of deoxygenated blood, contained 50 mM sodium dithionite. A 100 µm diameter Drummond micropipet (1/4 µL volume) was horizontally inserted into the drop, and the transit of the blood was recorded as it was drawn into the capillary micropipet. The position of the blood is readily observed by the unaided eye, however, and the recording was simply done for our own ancillary analysis of the flow, but is not essential to the method. All deoxygenated samples had their state of deoxygenation verified by a spectrophotometer described elsewhere, immediately following the transit measurement.

## Results

Typical data is shown in Fig. 1, which displays oxygenated and deoxygenated sickle blood in transit through the capillary. The flow of AA blood is the same for both oxygenated and deoxygenated blood. The transit for oxygenated blood from a Sickle-cell Anemia patient is somewhat faster than AA because the anemia reduces the hematocrit (HCT, the volume taken up by the red cells), and thus makes the solution less viscous. Deoxygenated SS blood not only flows substantially slower, but also eventually forms a dense region of cells behind the meniscus, that causes it to progress even slower. Because the analysis of horizontal flow is simpler than vertical flow, we collected the majority of this data in the horizontal geometry. However, vertical experiments revealed similar phenomena, viz. the slowed transit of deoxygenated sickle blood and the formation of a dense cap behind the meniscus. The vertical transits were also longer than the horizontal (e.g. ∼15 s vertical vs 10 s horizontal).

**Fig. 1.**
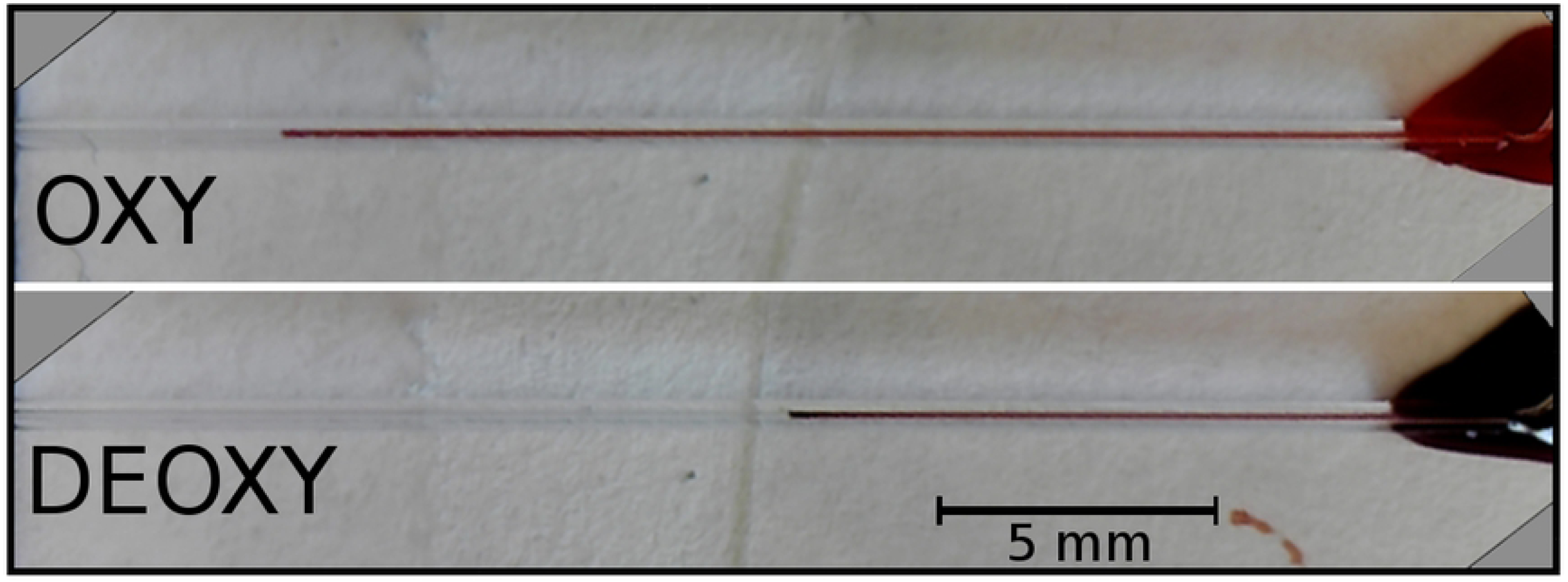
Movement of blood drawn into a horizontal capillary after 3 seconds, viewed from above. Capillaries were standard ¼ µL Drummond microcaps, approximately 3 cm long. The oxygenated blood on the left has clearly gone farther in the same time than the deoxygenated blood on the right. A drop of blood is placed on an 18 mm square coverslip into which a capillary has been inserted. In addition to the blood imbibed into the capillary itself, some blood runs along the space between the capillary and the coverslip, accounting for the odd shape of the source droplet.

Fig 2 shows how the time for cell transits varies for a range of genotypes. Each column in the figure displays the data for one individual, with each point designating a single measurement. The oxygenated blood behaves very consistently among the replications, as well as between individuals, regardless of genotype. As mentioned above, a degree of systematic variation is expected since the HCT varies between the genotypes. The dashed horizontal line shows the average oxygenated cell transit time for AA cells. The various sickle cell genotypes, including the patients being treated with hydroxyurea (HU group), have deoxygenated transit times which clearly exceed AA, and also showing greater variability.

**Fig. 2.**
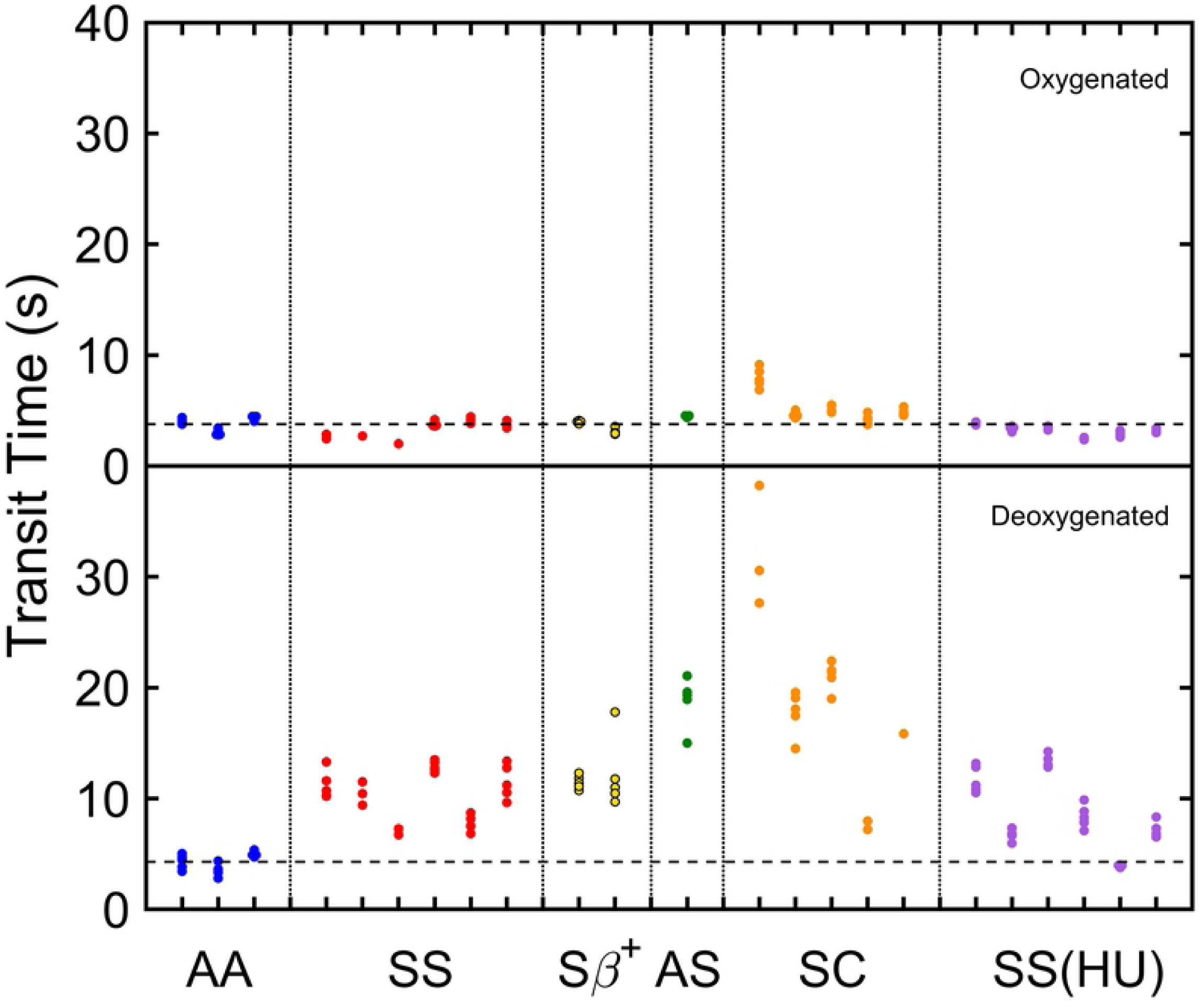
Time for cell transits along a distance of 24 mm in ¼ µL glass capillaries, for the genotypes studied. HU indicates patients being treated with hydroxyurea. Each column in the figure displays the data for one individual, and each point is a single measurement. The various sickle cell genotypes, including the HU group, clearly differ from AA, showing greater variability, and longer transits overall.

Another way to display the data of Fig 2 is to designate x and y axes as the transit time of the oxygenated and deoxygenated blood, respectively, so that each experiment becomes a point on that plane as shown in Fig 3. Each genotypic cluster (oxygenated and deoxygenated times) was characterized by a mean and standard deviation, the latter effectively displaying the patient-to-patient variation, since there was much less difference between individual replication experiments than seen between different patients. These standard deviations generate the ellipses in Fig. 3, centered on the mean value we obtained for each genotype. An elliptical zone of paired transit times thus emerged as a characteristic fingerprint of the genotype. For AS blood, only one patient was studied, and that ellipse is taken as a representative value from the other data sets.

**Fig. 3.**
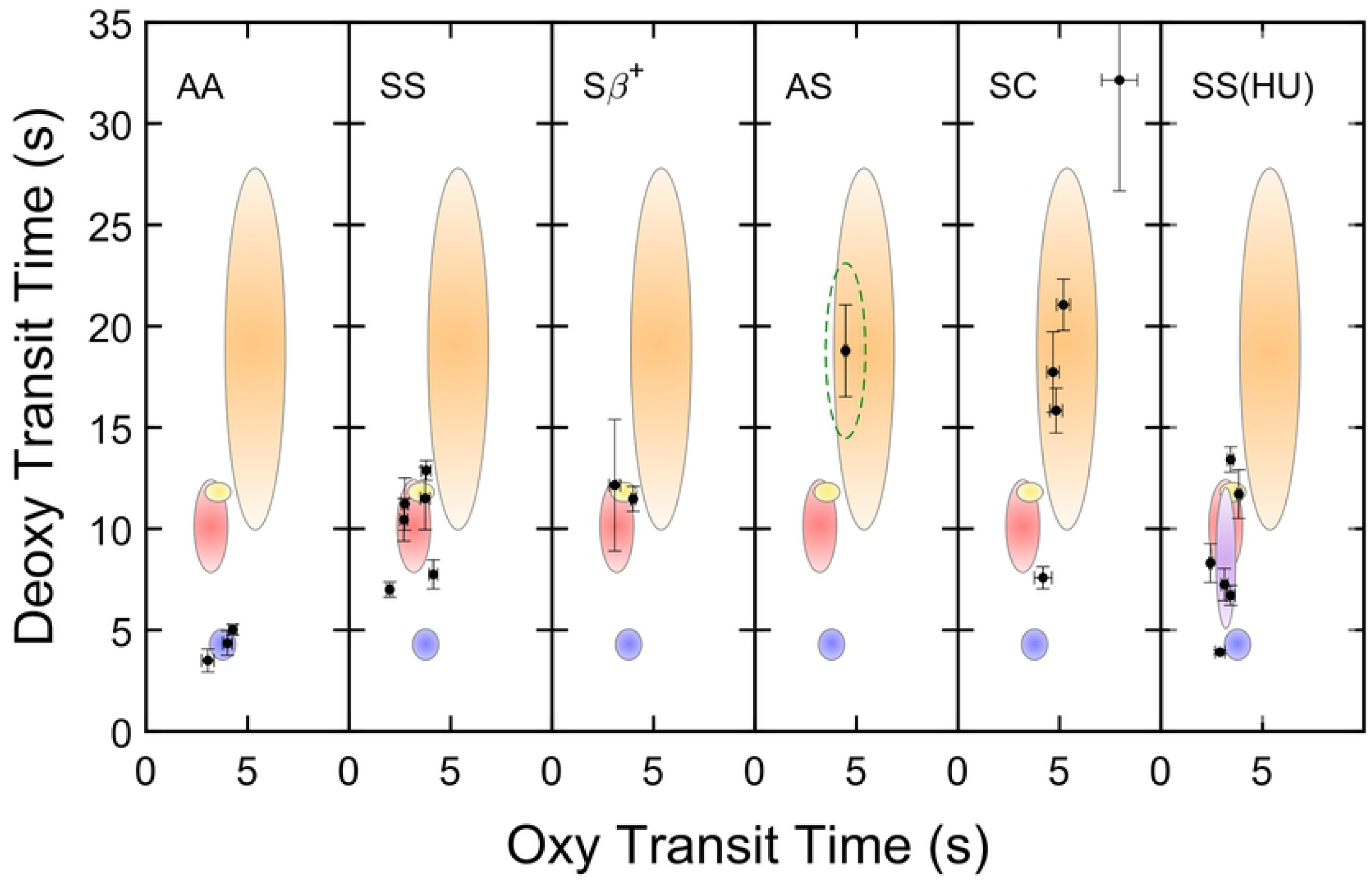
Transit time profiles. Data points from Figure 2 are displayed, with error bars and means that represent the dispersion and mean of the data from Fig. 2. An elliptical region has been constructed with center at the mean, and semi-major axes representing the standard deviation of that data, for each of the genotypes. Since only one patient was measured for AS, only a dashed ellipse is shown, approximated with axes representative of the group of all genotypes. Each point shown designates data for a single patient, and error bars of the point show the replication error for the particular patient. AA blood is well distinguished. For patients on the drug hydroxyurea (HU), the data spans the range from the SS to AA, suggesting the utility of this method in rapidly and inexpensively monitoring the response to the drug. It is known that not all patients respond similarly to this drug, which we suspect is the major factor in the range of variation of the HU data.

The AA region in Fig. 3 is clearly distinct from the sickling disorders. The region for SS patients shows a significant overlap with that for patients with Sβ^+^ thalassemia, but it is distinct from AS and SC patients.

We can generate an estimate of the sensitivity and specificity of this method. To this end, we generated continuous distributions, which had the mean and standard deviations calculated from the discrete data we collected. (For AS, having only one patient we used that mean value but adapted the standard deviation We took standard deviations from the other populations and divided them by their corresponding mean to get percent deviations. We then averaged the oxygenated and the deoxygenated percentages. Finally we multiplied the AS mean by the averaged percent to get estimated population standard deviations. Since such a continuous distribution will extend into regions with negative transit times, which are physically meaningless, we limited the distributions to physically possible values, and adjusted the normalization of the distribution accordingly.

The sensitivity of the test for any genotype area is defined as the ratio of true positives to the sum of true positives and false negatives. The specificity of a genotype is defined as the ratio of true negatives to the sum of true negatives and false positives. True positives were calculated by integrating the distribution over the region where it exceeded all other distributions (i.e. over its dominant region), while the false negative was calculated integrating the distribution over all regions except for that where the distribution exceeded all others. For the true negatives and false positive, the remaining distributions other than the genotype in question were integrated outside or inside the dominant region, respectively. As confirmation of this procedure, we generated a large random set of data with a gaussian distribution, using the given means and standard deviations, and found the results computed agreed to within 2 percent. These results are shown in Table 1, which provides a quantitative measure for what Fig 2 and 3 show intuitively. AA is readily discerned compared with any sickle variant. The same method picks SS correctly about 2 out of 3 times, confusing it most often with Sβ^+^ thalassemia.

**Table 1.** Sensitivity and specificity.

|  | Sensitivity | Specificity |
| --- | --- | --- |
| AA | 99% | 98% |
| SS | 70% | 97% |
| SC | 57% | > 99% |
| AS | 77% | > 99% |
| Sβ<sup>+</sup> | 94% | 98% |

We have also measured blood from patients who are being treated with the drug Hydroxyurea (HU), one of the two drugs approved for sickle cell disease [14]. HU increases the production of Fetal Hb, which is incapable of polymerization [12], and modestly increases the HCT from 25 to 27%. [14] The final panel of Fig 3 shows the timing profile we measured in 6 sickle patients currently on HU. The profile extends from the top of SS toward that of AA patients. (We did not include HU patients in the specificity and sensitivity calculations since such patients, *ipso facto*, already have their genotype identified.)

Since the basis of this test is the difference in the viscosity of sickle blood, we also show how the viscosity varies for the various genotypes. Figure 4a is a “fingerprint” analogous to Fig. 3, with viscosity shown instead of transit time for deoxygenated vs oxygenated blood. As found for simple timing measurements, so too for viscosity: the AA genotype is well distinguished. If the deoxygenated viscosity is normalized by the oxygenated viscosity and the ratio is plotted, the result, in Fig 4b, shows remarkable similarity amongst the various genotypes. The factor of increase in viscosity for undeformable sickle cells is, within error, the same ratio as seen upon the rigidification of AA red cells, shown by Chien et al. [13] From this one may conclude that it is the transformation of flexible cells to rigid particles which dominates the ratio, irrespective of the shape the particles may have assumed when they lost their deformability.

**Fig. 4.**
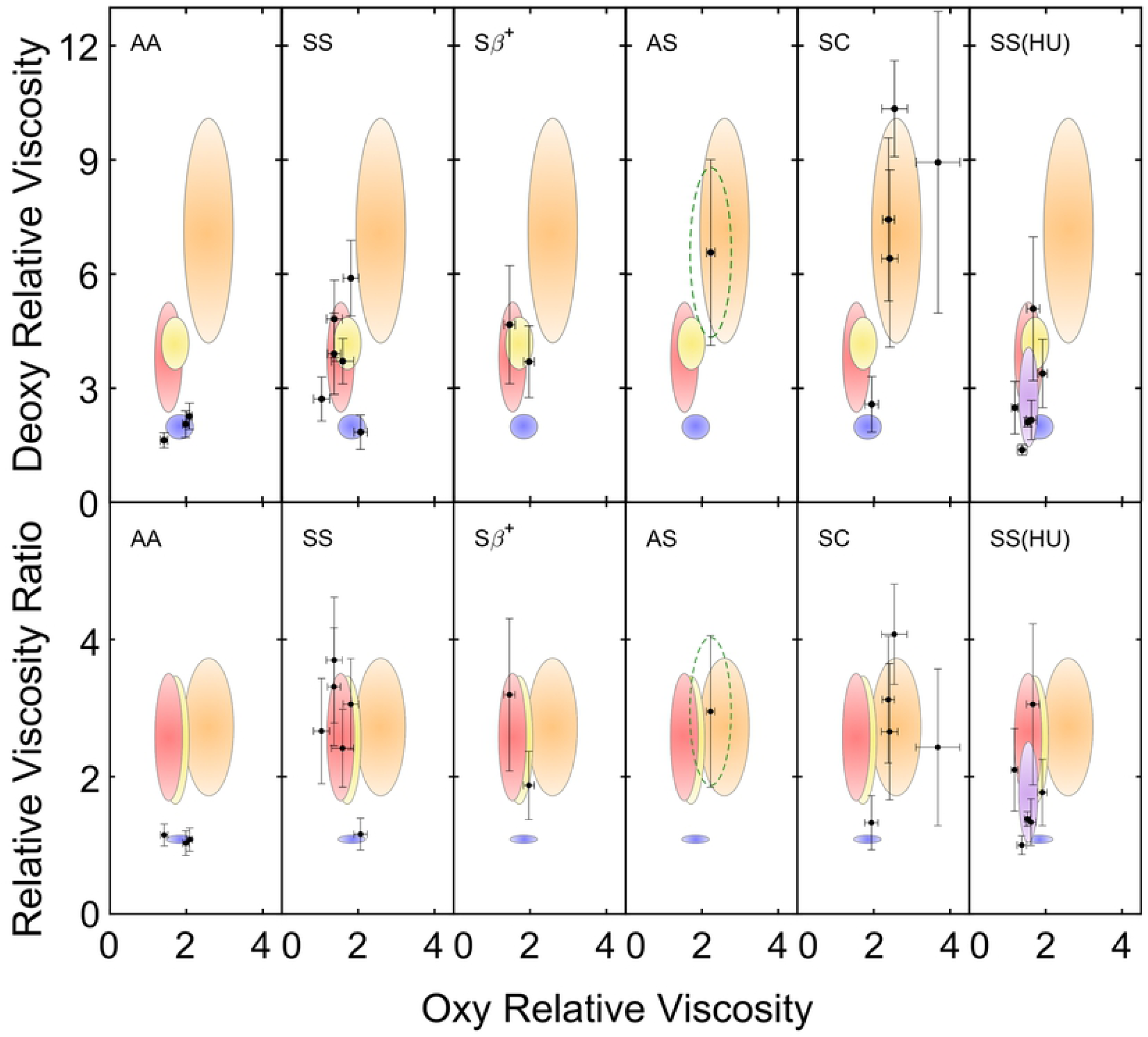
Viscosity and viscosity-ratio profiles. (a) Viscosity has been deduced from the data shown in Fig 2 and 3. Deoxygenated viscosity is plotted as a function of oxygenated viscosity for the various genotypes as well as patients on HU. As for Fig. 3, AA blood is well distinguished. (b) Ratios of deoxygenated to oxygenated viscosity, shown as a function of oxygenated viscosity. The ratio and its standard deviation are remarkably consistent among the sickle genotypes.

## Discussion

It is apparent that the sickling genotypes can clearly be distinguished from normal AA blood by the method shown here. A quantitative analysis gives a 98% true positive rate for distinguishing AA patients from those who possess the sickle gene. The other genotypes are distinguishable from one another to varying degrees, though all are distinct from AA blood. SS and Sβ^+^ thalassemia differ from AS and SC, primarily based on their hematocrit dependence. However, AS and SC are quite similar in this test, as are SS and Sβ^+^ thalassemia.

The use of paired oxy and deoxy values in Fig. 3 avoids a number of potentially confounding issues. For example, polycythemia vera generates high hematocrits. These would increase viscosity and thus slow the transit of blood through the test capillary. However, the concomitant increase in the viscosity of oxy *and* deoxy blood differs from the sickling disorders where only deoxygenated blood increases. In Fig 3, such a disorder of increased viscosity would displace the AA points diagonally upward, whereas SS displaces these points almost vertically (with a slight left shift). In a similar fashion, the presence of malarial parasites would increase viscosity of oxy and deoxy cells alike [15]. Moreover, the viscosity increase from malaria is also more modest than observed here. Finally, the same rationale applies to temperature effects. The viscosity of blood decreases about 30% as temperature is raised from 25°C to 45°C [15]. The temperature dependence is the same as that of water [16], and the temperature changes would again move data diagonally along the plot.

The use of a vertical geometry for the capillary insertion into the drop would serve to enhance resolution further. It is straightforward to show (cf paper 1, eq 1) that to traverse a given capillary length, deoxy blood takes longer than oxy blood by the same factor, independent of vertical or horizontal alignment. But, since vertical rise is intrinsically slower (thanks to gravity opposing capillarity), the differences are amplified in the vertical direction. Vertical insertion into a drop of blood would likely be more easily arranged for day to day clinical study.

The present data are taken by initial deoxygenation in a vial using sodium dithionite solution, followed by extraction of a 5 µL drop. However, that technique was devised for laboratory convenience, not field use, and it is not hard to imagine the development of pre-loaded vials, or other methods (such as using other dithionite-laden capillaries for deoxygenation) as a first step to expedite and simplify this procedure. Likewise, although we obtained detailed data with a small camera the same diagnostic value is readily obtained by measuring the transit time to some fixed distance, which can be done by visual observation and stopwatch timing (c.f. SI Fig. 1).

It is useful to compare this capillary test to the handful of other point of care tests. Four such tests have been published and have been reviewed in detail. [17].

In the first of those four tests, a drop of lysed, deoxygenated blood is placed on a paper and the diffusion of the deoxygenated hemoglobin is observed. [18-22] Polymerized HbS is relatively immobile compared to HbA, and AS has a mix of sickled and unsickled Hb with the result that these three genotypes can be distinguished, as with the metric shown here. The test requires the mixing of 20 µL of blood with sickledex solution, followed by a wait of about 35 minutes after application of the drops to the paper. Similar to our findings, AS and SC are difficult to distinguish, as are SS and Sβ^+^thalassemia.[22] In comparison to such a paper test, the capillary method has similar discrimination, is equally inexpensive, but is dramatically faster since the rise in the capillaries is complete in seconds, not minutes.

Two tests of greater accuracy use antibody-based methods. Both begin with cell lysis. One test using monoclonal antibodies [23] distinguishes AA, SS, AS and SC. This discrimination is superior to the capillary test, though the antibody mis-diagnoses Sβ^+^ thalassemia as AS. The test takes typically 20 minutes. A different immunoassay [24] successfully resolves the all the foregoing genotypes as well as Sβ^+^ thal, and succeeds in obtaining results in as little as 2 minutes after mixing the drawn blood drop with lysing agents, albeit at a cost of around $5 [17]. In comparison, the other two methods are ten times less expensive.

The fourth test described is a density-based system that does not require cell lysis. [25, 26] Aqueous Multiphase Systems (AMPS) are mixtures of polymers in water that form immiscible phases, and have been used with centrifugation with to reveal the presence of high-density cells that are a hall-mark of sickle cell disease. Of course, the method would not distinguish AS from AA, and also might be confounded by things such as liver disease or polycythemia vera that generate denser cells. This system, considered to take 12 minutes, requires the presence of a centrifuge, and given the inefficiency of running a solo tube, would likely mean tests are done in batches.

None of these methods match the combination of time resolution inherent in the capillary-based system (20 sec for the two measurements) and its low cost. The cost of the paper-based system might be comparable, but it takes far longer. The polyclonal antibody system is speedy but is still ∼6 times slower, and at $5 per test is far more costly than the capillary test. An optimal approach for diagnosis in resource-challenged regions might combine methods. Capillary flow as a pre-screen can rapidly and inexpensively sort AA patients from their sickle counterparts, for whom a follow-up test at higher cost might be appropriate. The value of such a strategy can be appreciated by considering places where the incidence is very high, such as Nigeria. There the S allele appears in about only 20% of the population [4], so that 80% of the tests performed in a general screen would reveal a non-S patient. For other nations with lower incidence, an even greater percentage of tests would be expected to have negative results. For this reason a rapid and inexpensive pre-screen could be very useful.

There is also information present in capillary tests that is simply not present in the antibody tests. As the final panel of Fig 3 shows, the behavior of blood for SS patients taking hydroxyurea stretches between the region of SS and that of AA, yet the different patients responded differently. Response of different patients to HU differs [17, 27]. The data of Fig 2 shows that different patients on HU have dramatically different deoxygenated blood transits. For some it is virtually identical to AA cells; for others it is much more like untreated SS blood. Having a rapid and inexpensive metric that could track such a response could be very useful.

An important question is whether the accuracy of the method can be improved. From Fig. 2 it is evident that the patient-to-patient variation is far greater than the variability of measurements on an individual patient. This is dramatically evident in the SC patients. Interestingly, when the measurement was repeated on the same AS patient after several months, the range of variation was virtually identical. We suspect the range of variability in SC and, by extension, AS, is a consequence of the complexity of the sickling process itself. The shapes of the sickled cells can range from the classic, extended eponymous sickle form to compact yet rigid cells that have been said to resemble potatoes. These shapes are exquisitely sensitive to the rate at which polymerization occurs, [28] which in turn depends on the cellular contents as well as the rate of deoxygenation. In particular, heterozygous conditions produce far greater distortions in the sickled cells than homozygous SS cells, a fact which is utilized in a current drug assay. [29] The differences within a given genotype might well be the consequence of the history of a given patient that shaped the particular cellular population at the time of observation. While this variable is viewed as confounding in the attempt to ascertain the presence of the S allele, it might actually provide unexpected insights, to the extent that the patients with longer transits might have a more severe course of the disease.

In this context, it is a welcome bonus that this test provides a snapshot of patient response to hydroxyurea. Response to this drug has proven to be quite variable. [27] The last panel of Fig 3 demonstrates this, as it is clear that the upper end of the distribution of deoxy transit times contains times equivalent to those of SS patients, suggesting the lack of any significant response to the drug. On the other hand, the low end of the distribution falls decidedly below that of SS patients, suggesting that these are the higher responders. (Due to the nature of our protocol, we had no other data that described the patients’ severity of clinical course.) Further testing is required to show whether this rapid point of care test might assist in providing feedback to patients and their physicians when this or other anti-sickling agents are in use.

## ACKNOWLEDGEMENTS

We thank Michelle Cahill and Dr. Nataly Apolonsky, of St. Christopher’s Hospital for Children, for provisions of samples and continued generous cooperation in support of this project, and Dr. William Eaton, of the Laboratory of Chemical Physics, NIDDKD, NIH, for providing us with valuable advice as well as discard AS blood.

